# Electrophysiological correlates of placebo hypoalgesia and nocebo hyperalgesia during the anticipation and delivery of pain

**DOI:** 10.64898/2026.11.17.766963

**Authors:** Antonia Baumgarten, Frederik Schlitt-Nguyen, Helena Hartmann, Katharina Schmidt, Michael Rose, Dagmar Timmann, Enzo Nio, Eline S. Kruithof

## Abstract

Positive and negative expectations can substantially shape pain and analgesic treatment outcomes, highlighting the need to elucidate neural mechanisms underlying placebo hypoalgesia and nocebo hyperalgesia. The primary aim of this feasibility study was to assess adaptations to an established placebo-nocebo conditioning pain paradigm in the context of electroencephalography (EEG). The secondary aim was to assess EEG correlates of placebo hypoalgesia and nocebo hyperalgesia during the anticipation and delivery of experimental heat pain in healthy volunteers. Placebo hypoalgesia and nocebo hyperalgesia were induced in a within-subjects design in 20 young adults through verbal instructions, a conditioning procedure and (sham) transcutaneous electrical nerve stimulation while EEG was recorded. Placebo and nocebo effects as well as positive and negative pain expectations were successfully established, demonstrating paradigm effectiveness. EEG results showed increased theta, alpha and beta power in the placebo compared to the control condition during pain delivery, and in the nocebo compared to the control condition during pain anticipation and delivery. Additionally, gamma-band power was increased in the nocebo compared to the placebo condition as a function of a higher perceived pain intensity. Taken together, feasibility of the EEG-compatible placebo-nocebo conditioning pain paradigm was demonstrated. Furthermore, our EEG results are in line with previous findings that, in the context of placebo hypoalgesia and nocebo hyperalgesia, link theta-, alpha- and beta-band activity to top-down processing, and gamma-band activity to expectation-modulated sensory processing.

## 1. Introduction

The experience of pain is influenced by expectations. Positive expectations can result in pain relief, which is known as placebo hypoalgesia, whereas negative expectations can result in pain worsening, which is known as nocebo hyperalgesia (Atlas & Wager, 2012). In an experimental setting, placebo hypoalgesia and nocebo hyperalgesia are often induced through a conditioning procedure combined with verbal instructions. Here, placebo and nocebo treatments are surreptitiously paired with a pain stimulus of low and high intensity, respectively, while a pain stimulus of medium intensity is applied in the control condition (Wager & Atlas, 2015). Placebo and nocebo effects can subsequently be tested by administering the same medium painful stimulus intensity in all conditions, and compare placebo and nocebo conditions with the control condition.

Electroencephalography (EEG) can shed light on neural processes associated with placebo hypoalgesia and nocebo hyperalgesia. Alpha- and beta-band activity have been linked to top-down pain processing (Ploner et al., 2017), which is based on cognitive, affective and motivational factors such as expectations (Tiemann et al., 2015), highlighting its relevance for placebo and nocebo effects. For example, in healthy volunteers, both a placebo and nocebo manipulation were shown to induce an increase in alpha power compared to a control condition and compared to high baseline pain (Albu & Meagher, 2016; Huneke et al., 2013; Thomaidou et al., 2021a). However, Tiemann et al. (2015) found no difference in alpha-band activity between placebo and control conditions. During pain anticipation, alpha and beta power have been linked to expected and perceived pain intensity, with both positive (Strube et al., 2021) and negative (Babiloni et al., 2003, 2006; Nickel et al., 2022; Tu et al., 2016) associations being reported. Conversely, gamma-band activity is suggested to mediate bottom-up pain processing (Ploner et al., 2017; Tiemann et al., 2015; Strube et al., 2021), which is driven by sensory properties of the stimulus, such as intensity (Tiemann et al., 2015). Furthermore, gamma-band activity is positively associated with subjective pain perception during pain anticipation and delivery (Gross et al., 2007; Tu et al., 2016; Taesler & Rose, 2016; Zhang et al., 2012). However, results on gamma-band activity during placebo and nocebo conditions go in mixed directions. That is, lower as well as higher gamma power have been reported in a nocebo condition compared to a control condition and high baseline pain (Thomaidou et al., 2021a; Mourkojannis et al., 2025), while no differences were found in a placebo compared to control condition (Tiemann et al., 2015).

The primary aim of this feasibility study was to assess adaptations made to an established placebo-nocebo conditioning pain paradigm (German Clinical Trial Register, 2023; Kunkel et al., 2025) in the context of EEG. Placebo hypoalgesia and nocebo hyperalgesia were induced through verbal instructions, a conditioning procedure and (sham) transcutaneous electrical nerve stimulation (TENS). Specifically, TENS was framed as a method of frequency-dependent pain modulation. Several notable changes were made to the paradigm. First, the heat stimulus was substantially shortened from 20 to 2 s and delivered with a steep heating rate (70 °C/s). This provides a well-defined onset to which EEG activity can be time-locked, as well as short and comparable epochs for event-related analysis rather than a long period over which pain perception can change. Second, the shorter heat stimulus duration allowed us to increase the number of trials in the test phase (i.e., in which placebo and nocebo effects are tested) to 120 while keeping session length and cumulative heat exposure acceptable. Third, expected pain intensity ratings were implemented in the first 90 test phase trials instead of at four time points. As these changes alter previous versions of the paradigm substantially, it was unclear whether placebo and nocebo effects would be preserved. For instance, a brief heat stimulus may produce weaker placebo and nocebo effects than a 20 s stimulus. Furthermore, across 120 test phase trials these effects may decline before enough data could be acquired to perform an EEG analysis that has sufficient power. In addition, rating expected pain intensity on a trial-by-trial basis may itself alter placebo and nocebo effects, for instance by keeping expectations explicitly active and slowing their reduction (Jepma et al., 2018).

The secondary aim of the study was to assess EEG correlates of placebo hypoalgesia and nocebo hyperalgesia during the anticipation and delivery of experimental heat pain in healthy adult volunteers. Based on the current literature, we hypothesized differences in alpha and beta power in the placebo and nocebo compared to the control condition during the anticipation and delivery of experimental heat pain. Furthermore, we hypothesized that gamma power would be increased during pain anticipation and delivery as a function of a higher perceived pain intensity between conditions (i.e., nocebo > control > placebo).

## 2. Methods

### 2.1 Participants

Twenty-two healthy adult volunteers participated in this within-subjects study. Participants were recruited via the social media channels of the Department of Neurology of the University Hospital Essen, a neighborhood website (nebenan.de) as well as via the websites of the student councils. All participants were 18-35 years old, right-handed or ambidextrous, reported normal or corrected-to-normal vision and had sufficient knowledge of German language to follow the instructions. Exclusion criteria were red-green color blindness, chronic diseases (including chronic pain), regular medication or drug use (including nicotine), neurological, psychiatric and internal diseases, hormonal contraception, pregnancy, breastfeeding, acute skin damage, dermatological diseases and a surgical procedure under general anesthesia within the last six months. Participants did not take part in other pain studies within the last 6 months. They were requested to refrain from alcohol and drugs in the 24 hours before the experiment and to refrain from caffeine on the day of the experiment. Two participants were excluded from the analyses, one due to technical issues during the experiment and one because individual differential heat stimulus temperatures could not reliably be determined, leading to a final sample of 20 participants (19 right-handed and 1 ambidextrous, 10 females, age (*M* ± *SD*) = 25.55 ± 4.21 years, age range = 18-35 years). The sample size was not determined a priori in a power analysis, as the study was exploratory. All participants gave written consent. Participants received a monetary compensation of 15 EUR per hour. The study was approved by the local ethics committee of University Hospital Essen and performed in accordance with the Declaration of Helsinki, version 2000 (World Medical Association, 2000).

### 2.2 Placebo-nocebo conditioning pain paradigm

The placebo-nocebo conditioning pain paradigm was used to examine the effect of experimentally induced pain-related expectations on pain perception. Expectations of pain relief (placebo condition), no change in pain (control condition) and pain worsening (nocebo condition) were induced through verbal instructions, a conditioning procedure and (sham) transcutaneous electrical nerve stimulation (TENS). More specifically, participants were led to believe that (sham) TENS modulates their heat pain perception depending on its frequency. They were told that different colored crosses on the screen indicated which frequency was applied. At the beginning of the conditioning phase, participants were informed that the green cross signaled low frequency electrical stimulation which relieves pain, the yellow cross signaled middle frequency electrical stimulation which does not have an effect on pain perception, and the red cross signaled high frequency electrical stimulation which worsens pain. Participants were told that differences between these TENS frequencies could not be perceived. Unbeknownst to the participant, heat stimulus temperatures were adjusted in each condition (placebo, control, nocebo) during the conditioning phase. That is, placebo, control and nocebo cues (i.e., the colored crosses) were paired with a heat stimulus temperature corresponding to an individually calibrated pain intensity of 40, 60 and 80 on a 0-100 visual analogue scale (VAS), respectively (0 = ‘not painful at all’, 100 = ‘unbearably painful’; **Figure 1A**). In the subsequent test phase, however, placebo, control and nocebo cues were all paired with a heat stimulus of the same temperature, corresponding to an individually calibrated pain intensity of 60 on the 0-100 VAS (**Figure 1B**). The conditioning phase consisted of 30 trials (10 per condition). The test phase consisted of 120 trials (40 per condition), distributed across 4 blocks of 30 trials each (10 per condition). The order of conditions was pseudo-randomized in both the conditioning and test phases. Before the conditioning phase, participants performed three practice trials, one for each condition, to familiarize themselves with the trial structure.

**Figure 1.**
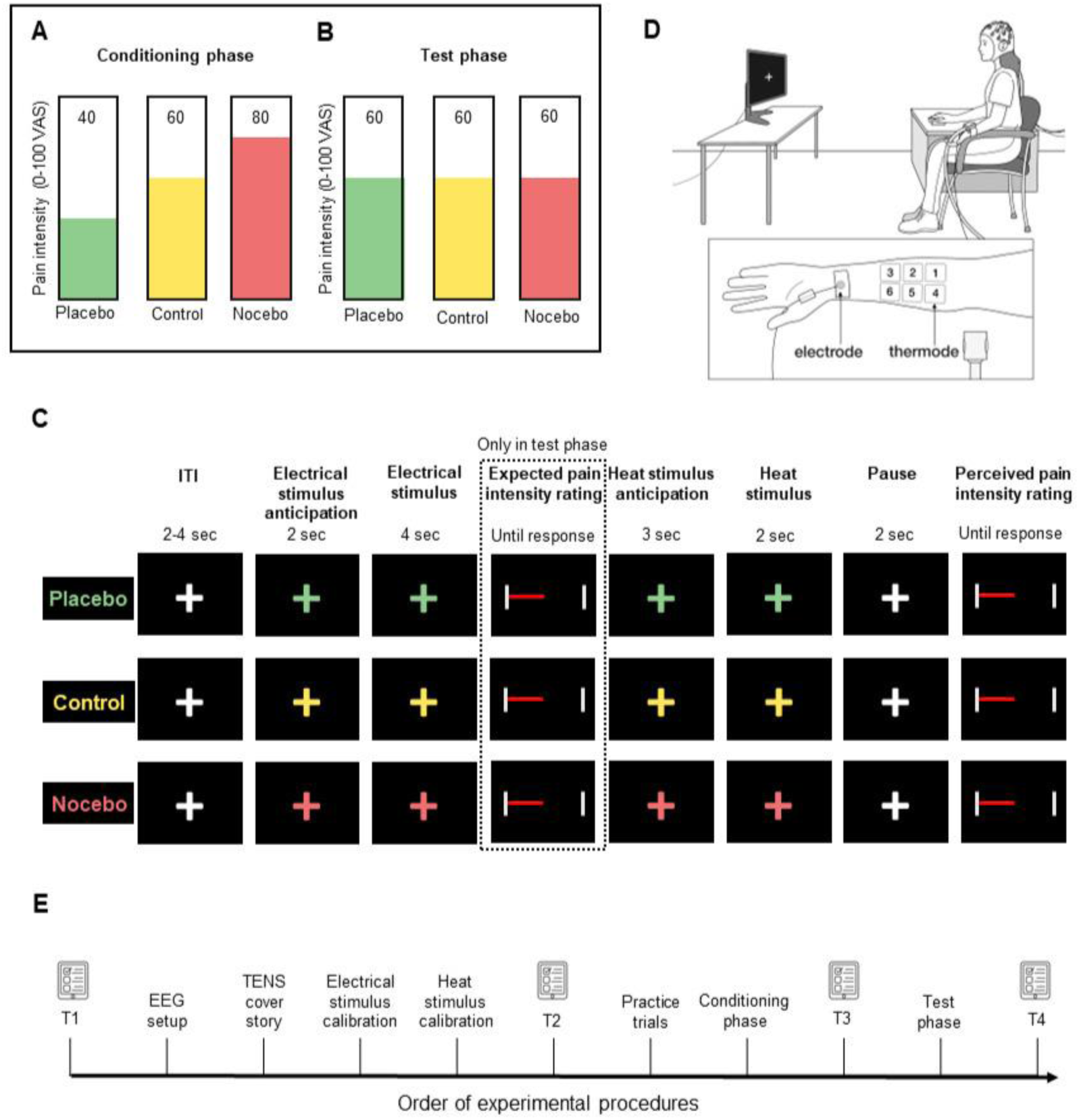
Placebo-nocebo conditioning pain paradigm and procedures. A) Study design in the conditioning phase. In the placebo, control and nocebo conditions, heat stimuli with temperatures corresponding to an individually calibrated pain intensity of 40, 60 and 80 on a 0-100 VAS, respectively, were applied. B) Study design in the test phase. In the placebo, control and nocebo conditions, heat stimuli of the same temperature, corresponding to an individually calibrated pain intensity of 60 on a 0-100 VAS were applied. C) Event sequence of a trial. After a jittered intertrial interval, a colored cue signaled the upcoming electrical stimulus (electrical stimulus anticipation), followed by the electrical stimulus itself. In the test phase only, participants next rated their expected pain intensity of the upcoming heat stimulus on the 0-100 VAS. Subsequently, the colored cue indicated the upcoming heat stimulus (heat stimulus anticipation), after which the heat stimulus was applied. Finally, after a pause, participants rated their perceived pain intensity of the heat stimulus on the 0-100 VAS. D) Experimental setup and positioning of the electrode and thermode on the left volar forearm. The thermode was positioned at 6 different locations during the experiment. E) Order of experimental procedures. T1, T2, T3 and T4 denote the four time points at which psychological questionnaires were administered: T1 (baseline), T2 (before the practice trials), T3 (before the test phase), T4 (after the test phase). EEG = electroencephalography, ITI = intertrial interval, TENS = transcutaneous electrical nerve stimulation, VAS = visual analogue scale.

The placebo-nocebo conditioning pain paradigm was adapted from Hartmann and colleagues (German Clinical Trial Register, 2023) and Kunkel et al. (2025) and performed in Presentation software (version 25.0, Neurobehavioral Systems, Inc., Berkeley, CA). **Figure 1C** shows the event sequence of a trial. Some modifications were made to the paradigm for compatibility with EEG. First, the duration of heat stimulus was shortened from 20 to 2 sec. Second, the number of trials in the test phase was increased to 120 trials. Third, the number of trials in the conditioning phase was reduced from 36 to 30 (10 per condition), in line with the number of trials per test phase block. Fourth, per-trial expected pain intensity ratings were implemented, as explained in the next paragraph. Fifth, the heat stimulus anticipation time window was shortened to a fixed duration of 3 sec. Furthermore, additional (non-EEG-related) modifications were made to the paradigm. First, event durations (i.e., intertrial interval, electrical stimulus anticipation) were adjusted. Second, the reaction time task at the beginning of the trial was removed.

In the conditioning phase as well as in test phase blocks 1-3, participants rated the perceived pain intensity after each heat stimulus on a VAS ranging from 0 (‘not painful at all’) to 100 (‘unbearably painful’). In test phase blocks 1-3, participants additionally rated the expected pain intensity of the heat stimulus on a VAS ranging from 0 (‘not painful at all’) to 100 (‘unbearably painful’). For both ratings, the cursor was located on a random position on the scale, which participants could move to adjust and confirm their rating using a button box without time restrictions. Participants were instructed to respond as intuitively as possible. Test phase block 4 did not include per-trial perceived and expected pain intensity ratings. This block was implemented for a future study in patients with cerebellar degeneration to control for the possible influence of motor performance deficits on EEG activity. Therefore, data of this block were not used for analyses.

### 2.3 Electrical stimulation and calibration

Electrical stimuli were delivered via a 6.5 mm concentric surface electrode (WASP electrode, Specialty Developments, Bexley, UK) connected to a Digitimer DS7A constant current stimulator (Welwyn Garden City, UK). The electrode was attached to the left volar forearm, approximately 2.5 cm above the wrist (**Figure 1D**). Stimulation consisted of a 4-sec train of 0.1-ms pulses with a 30-ms interpulse interval.

The intensity of the electrical stimulus was determined individually in a calibration procedure (German Clinical Trial Register, 2023; Kunkel et al., 2025). After determining the detection threshold, four 4-sec electrical stimuli were administered and rated on a VAS ranging from 0 (‘not perceivable) to 100 (‘unbearably painful’). The intensity that participants rated as approximately 33, corresponding to ‘perceivable but not painful’ was the intensity used during the conditioning and test phases. If this target intensity was not achieved with 4 stimuli, the rating procedure was repeated until the stimulus intensity that corresponded to a VAS of approximately 33 could be determined.

### 2.4 Heat stimulation and calibration

Heat stimuli of 2 sec were delivered from a baseline of 35 °C at a heating rate of 70 °C/s and cooling down rate of 40 °C/s using an advanced thermal stimulator (PATHWAY system, model CHEPS, 27 mm diameter, Medoc Ltd., 2005).

The intensity of the heat stimuli was individually calibrated (German Clinical Trial Register, 2023; Kunkel et al., 2025). First, the heat pain threshold (HPT) was determined according to the method of limits. The subsequent calibration procedure consisted of 21 heat stimuli with varying temperatures around the HPT (-1 °C to 3.5°C), which participants rated on a VAS ranging from 0 (‘not painful at all’) to 100 (‘unbearably painful’). These ratings (except the first, which was applied to habituate the skin to the stimulation) were entered into a linear regression to estimate average temperatures corresponding to a pain intensity of 40, 60 and 80 on the VAS. Each of these temperatures was subsequently applied twice in a short test in a pseudo-randomized order, to verify the final temperatures. If necessary, final temperatures corresponding to a pain intensity of 40, 60 and 80 on the VAS were adjusted. For safety reasons, a maximum temperature of 50.5°C was set. To prevent sensitization or habituation, the thermode was positioned at 6 different locations during the experiment (**Figure 1D**). The thermode was repositioned before the conditioning phase and before each test phase block. The conditioning phase and each test phase block began with a reference trial, in which a gray cross was paired with a heat stimulus administered at the control condition intensity (i.e., pain intensity of 60 on the VAS). Initial thermode position was randomized across participants and then consecutively changed in ascending order. Furthermore, initial thermode position was randomized with the order of conditions in the conditioning and test phases across participants.

### 2.5 Electroencephalography (EEG) recording

EEG was recorded from 59 active Ag/AgCl electrodes (Fp1, Fp2, AF7, AF3, AFz, AF4, AF8, F7, F5, F3, F1, Fz, F2, F4, F6, F8, FT7, FC5, FC3, FC1, FCz, FC2, FC4, FC6, FT8, T7, C5, C3, C1, Cz, C2, C4, C6, T8, TP7, CP5, CP3, CP1, CPz, CP2, CP4, CP6, TP8, P7, P5, P3, P1, Pz, P2, P4, P6, P8, PO7, PO3, PO4, PO8, O1, Oz, O2) positioned according to the international 10-10 system. POz served as the reference and Fpz as the ground electrode. The four remaining electrodes (TP9, TP10, FT9 and FT10) were used to record a vertical and horizontal electrooculography (EOG) and an electromyography (EMG). Two electrodes were fixed below the outer canthi of each eye as horizontal electrooculogram (hEOG) and one electrode below the left eye as vertical electrooculogram (vEOG). The EMG electrode was positioned above the right descending part of the trapezius muscle. Data were sampled at 1000 Hz and amplified with two BrainAmp amplifiers (Brain Products GmbH, Gilching, Germany). Impedance was kept below 15 kΩ. EEG was recorded during the conditioning and test phases.

### 2.6 Psychological questionnaires

Psychological questionnaires were administered as part of the transregional collaborative research center (CRC/TRR) 289 ‘Treatment Expectation’ (https://treatment-expectation.de/). The following questionnaires were administered: Big Five Inventory-10 (BFI-10), Fear of Pain Questionnaire III (FPQ-III), Generic rating scale for previous treatment experiences, treatment expectations, and treatment effects (GEEE), Interoceptive Sensitivity and Attention Questionnaire (ISAQ), Pain Catastrophizing Scale (PCS), Perceived Stress Scale (PSS-10), Somatosensory Amplification Scale (SSAS), State-Trait Anxiety-Depression Inventory (STADI), Warmth & competence. Questionnaires were administered at four time points: T1 (baseline), T2 (before the practice trials), T3 (before the test phase), T4 (after the test phase). The GEEE was administered at all time points (i.e., previous treatment experiences subscale at T1, treatment expectations subscale at T2 and T3, and experienced treatment effects subscale at T3 and T4). The trait anxiety and depression subscales of the STADI were administered at time point T1 and the state anxiety and depression subscales of the STADI were administered at time points T2-T4. Warmth & competence was administered at time point T4. The other questionnaires were administered at time point T1 only. Details about the psychological questionnaires are provided in **Supplementary Table 1**, and descriptive statistics of the psychological questionnaires are provided in **Supplementary Table 2.**

### 2.7 Procedure

Prior to study enrollment, participants received the study information and were screened for eligibility via telephone and e-mail. Participants were told that the aim of the study was to investigate the effect of TENS on individual heat pain sensitivity. After the opportunity to ask questions at the beginning of the experiment, participants signed the consent form. Next, the questionnaires of time point T1 were administered (i.e., BFI-10, FPQ-III, GEEE previous treatment experiences subscale, ISAQ, PCS, PSS-10, SSAS, STADI state anxiety and depression subscales), after which the participant was set up for EEG. Participants were then explained the TENS cover story, which was followed by calibration of the electrical and heat stimulus. During the experiment, participants were seated in a chair in front of a computer screen, with the left arm positioned on an armrest and the right arm positioned on a cabinet to control the button box with the right hand (**Figure 1D**). Before the practice trials of the conditioning phase, participants completed the questionnaires of time point T2 (i.e., GEEE treatment expectations subscale and the STADI state anxiety and depression subscales). Before the test phase, participants completed the questionnaires of time point T3 (i.e., GEEE treatment expectations and experienced treatment effects subscales, STADI state anxiety and depression subscales). After the test phase, participants completed the questionnaires of time point T4 (i.e., GEEE experienced treatment effects scale, STADI state anxiety and depression scales, Warmth & competence). **Figure 1E** shows the order of the experimental procedures.

### 2.8 Data analysis

#### 2.8.1 Placebo and nocebo effects and positive and negative expectations

Differences in perceived and expected pain intensity ratings between conditions and across test phase blocks were analyzed using repeated measures ANOVAs based on participant-level means. Perceived pain intensity ratings were analyzed separately for the conditioning phase and the test phase. Expected pain intensity ratings were analyzed for the test phase only, as these were not obtained during the conditioning phase. Repeated measures ANOVAs included the within-subjects factor *condition* (placebo, control, nocebo) and, in the test phase, additionally the within-subjects factor *test phase block* (Test 1, Test 2, Test 3). Note that during the conditioning phase, the within-subjects factor *condition* reflected the different stimulation temperatures administered (40, 60, and 80 on the 0-100 VAS for the placebo, control, and nocebo conditions, respectively). Significant main effects and interactions were followed up with Holm-corrected post hoc dependent samples *t*-tests. To examine the presence of placebo and nocebo effects (based on perceived pain intensity ratings) as well as positive and negative expectations (based on expected pain intensity ratings) in the test phase, main effects of *condition* were followed up by comparing the placebo with the control and the nocebo with the control condition in dependent samples *t*-tests. Additionally, dependent samples *t*-tests were performed to examine placebo and nocebo effects within each test phase block (Test 1, Test 2, Test 3). Repeated measures ANOVA assumptions of normality and sphericity were evaluated. Normality checks were treated as diagnostic rather than strict criteria for selecting alternative tests. As Mauchly’s test indicated violations of sphericity, Greenhouse-Geisser-corrected *p*-values are reported for effects involving repeated measures factors with more than two levels.

For the test phase, the placebo effect (based on perceived pain intensity) and positive expectations (based on expected pain intensity) were both defined as the control - placebo condition rating difference. The nocebo effect (based on perceived pain intensity) and negative expectations (based on expected pain intensity) were both defined as the nocebo - control condition rating difference. Perceived and expected pain intensity ratings were pooled across test phase blocks (Test 1, Test 2, and Test 3). As condition differences during the conditioning phase reflect different stimulation temperatures rather than expectation-driven and conditioned effects, conditioning phase differences in perceived pain intensity ratings are referred to as pain relief (control - placebo) and pain worsening (nocebo - control). Holm-corrected dependent samples *t*-tests were conducted to examine differences in magnitude between pain relief and pain worsening during the conditioning phase, and placebo and nocebo effects as well as positive and negative expectations during the test phase.

All analyses were performed in MATLAB R2024b (The MathWorks). The alpha level of significance was set to 0.05 (two-tailed) throughout.

#### 2.8.2 EEG

EEG data were preprocessed and analyzed using the FieldTrip toolbox version 20250523 (Oostenveld et al., 2011). Only the data of the test phase were used for analysis. First, EEG data were epoched and time-locked to heat stimulus onset. Each epoch ranged from 3.5 sec before to 2.5 sec after heat stimulus onset, which included the pain anticipation time window (3 sec before heat stimulus onset) and the pain delivery time window (2 sec after heat stimulus onset). Epochs were larger than the time window of interest to reduce edge artifacts in the time-frequency decomposition later on. After epoching, the data were band-pass filtered (4-128 Hz, Butterworth fourth order), centered (subtraction of the temporal mean) and detrended. Epochs were then visually inspected, and trials that contained large movement or technical artifacts were removed. On average, 1.46 % of the trials (*SD* = 2.81 %, range = 0-9.17 %) were removed per participant. Subsequently, automatic artifact rejection was applied to exclude epochs with signals exceeding +/- 200 µV. On average, 2.39 % of the trials (*SD* = 5.65 %, range = 0-24.77 %) were excluded per participant. An Independent Component Analysis (ICA) was then performed using a logistic Infomax algorithm implemented in EEGLAB version 2025.0.0 to identify and discard eye movement and muscle activity artifacts. These artifacts were additionally identified via correlations of the ICA components with the EOG and neck muscle channels. On average, 6.05 (*SD* = 2.36, range = 2-12) out of the 59 components were rejected per participant. After the ICA, the implicit reference POz was added and the data were re-referenced to the average of all channels, after which the implicit reference was removed. The data were then segmented into blocks with (blocks 1-3) and without (block 4) perceived and expected pain intensity ratings as well as in conditions (placebo, control, nocebo). Time-frequency decomposition was performed for 21 logarithmically scaled frequencies between 4-128 Hz (0.25-octave increments) using a sliding Hanning window Fourier transform with a window length of 500 ms and a step size of 50 ms. A shared expectation condition, reflecting activity shared by the placebo and nocebo conditions, was derived by averaging placebo and nocebo time-frequency representations for each participant.

To examine how time-frequency patterns during pain anticipation and delivery differ between conditions (placebo, control, nocebo), dependent samples *t*-tests were performed comparing pairs of conditions in the time window from 3 sec before to 2 sec after heat stimulus onset. Additionally, to examine overlap in time-frequency patterns between the placebo and nocebo condition, a dependent samples *t*-test was performed which compared the shared expectation condition (i.e., activity shared by the placebo and nocebo conditions) with the control condition. To test the hypothesis that gamma power relates to perceived pain intensity, linear regressions were performed to examine how differences in EEG power between pairs of conditions vary as a function of differences in perceived pain intensity ratings between the respective conditions. For each participant, differences in mean perceived pain intensity ratings between conditions (control - placebo, nocebo - control and nocebo - placebo) were computed. Here, a larger value indicates a larger difference in perceived pain intensity between the two conditions. The mean perceived pain intensity difference rating was used in the linear regression as predictor of the corresponding time-frequency representation difference map. Main analyses were restricted to test phase blocks 1-3 in which perceived and expected pain intensity ratings were acquired.

Analyses were corrected for multiple testing using non-parametric cluster-based permutation tests (Maris & Oostenveld, 2007). Samples with a statistic that met the cluster-forming threshold of *p* ≤ 0.05 were clustered based on spatial (i.e., at least one significant neighbor channel required for the sample to be included), temporal and spectral adjacency. Neighbors were defined according to a template of the FieldTrip toolbox that corresponded to the EEG montage used. Cluster-level statistics were calculated as the sum of *t*-values within each cluster. Statistical significance was assessed for *α* = 0.05 (two-sided) by performing a Monte Carlo permutation procedure with 1000 permutations, which created a null distribution by randomly shuffling condition labels within participants and computing the cluster-level statistics for each permutation. Observed cluster statistics were then tested against this null distribution.

## 3. Results

### 3.1 Behavioral results

#### 3.1.1 Pain relief and pain worsening during the conditioning phase

A repeated measures ANOVA for perceived pain intensity ratings during the conditioning phase revealed a significant main effect of *condition* (*F*(2,38) = 185.65, *p* < 0.001). Post hoc *t*-tests showed significant pain relief indicated by lower perceived pain intensity ratings in the placebo compared to the control condition (mean difference = -23.59; *t*(19) = -9.18, *p* < 0.001, 95% CI [-28.97, -18.21]) and significant pain worsening indicated by higher perceived pain intensity ratings in the nocebo compared to the control condition (mean difference = 29.76; *t*(19) = 12.31, *p* < 0.001, 95% CI [24.70, 34.82]; **Figure 2A**). A direct comparison between pain relief and pain worsening showed no significant difference (*t*(19)= -1.64, *p* = 0.118, 95% CI [-14.06, 1.72]; **Figure 3A**).

**Figure 2.**
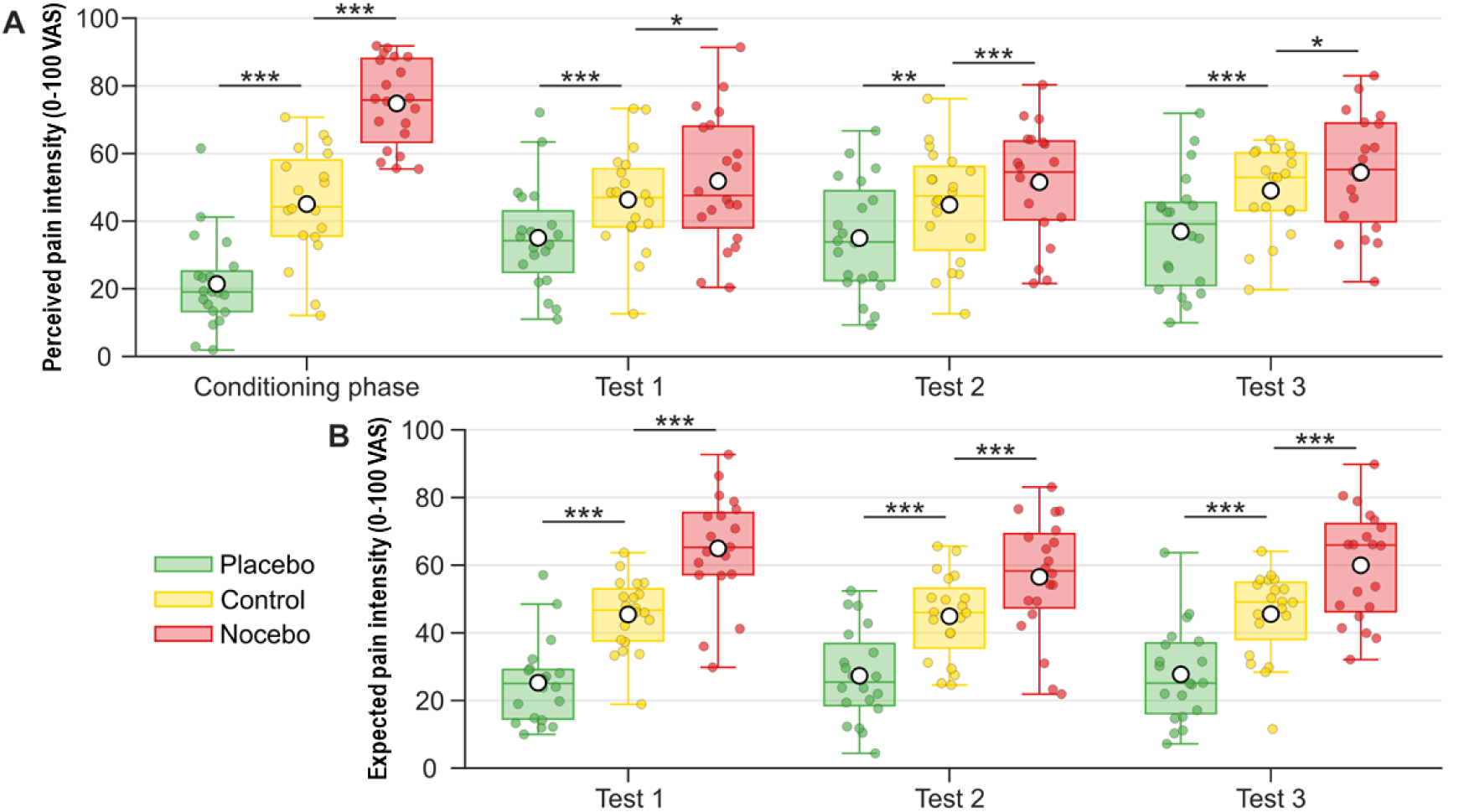
Mean perceived pain intensity (A) and expected pain intensity (B) for the placebo, control, and nocebo conditions during the conditioning phase (A only) and the three test phase blocks (i.e., Test 1-3). Boxplots display the median and interquartile range. Colored circles indicate individual participant means. White circles indicate group means. Asterisks indicate post hoc comparisons against the control condition (\**p* < 0.05, \*\**p* < 0.01, \*\*\**p* < 0.001). VAS = visual analogue scale.

**Figure 3.**
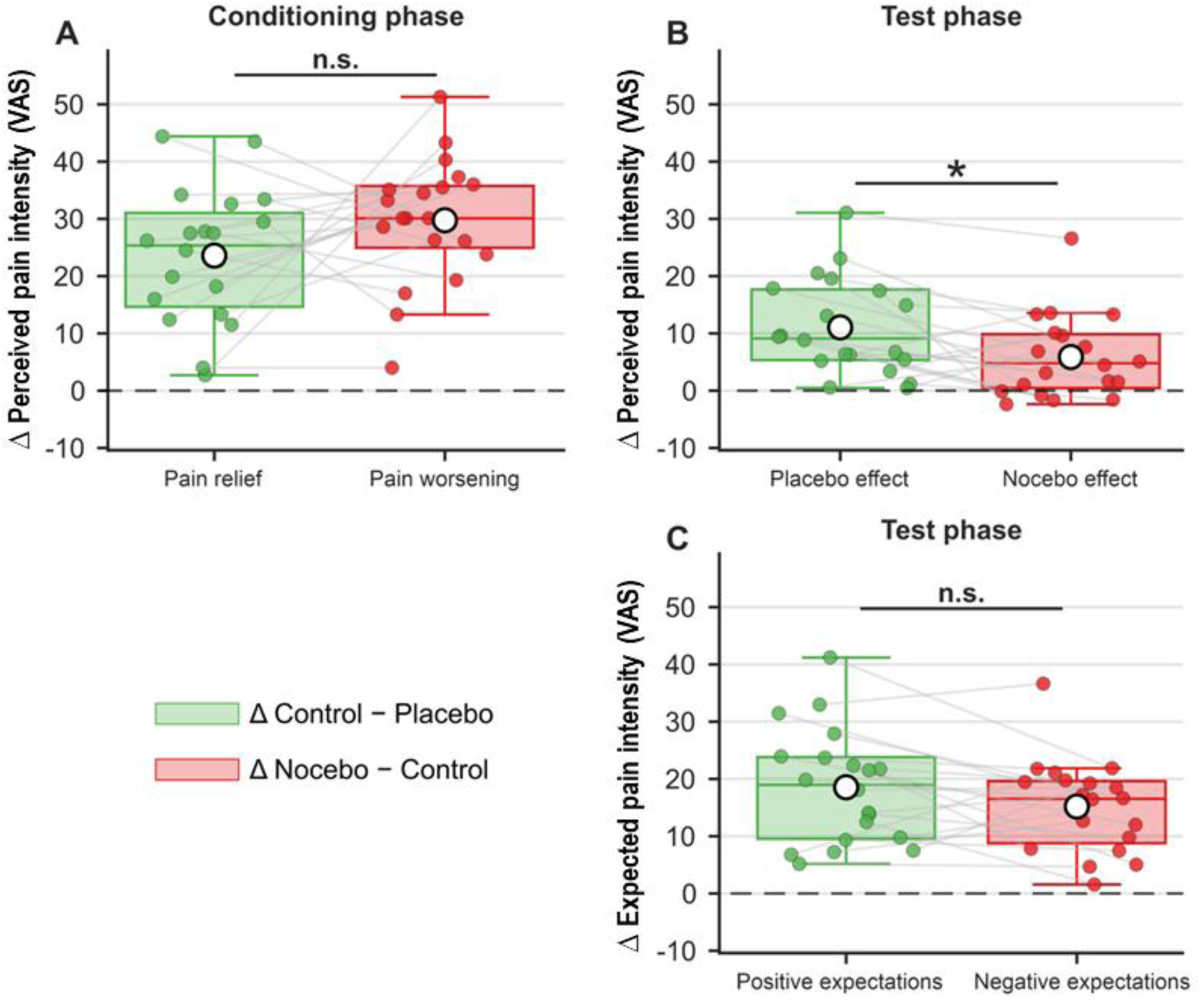
(A) Pain relief (Δ control – placebo: green) and pain worsening (Δ nocebo – control: red) based on perceived pain intensity ratings in the conditioning phase. (B) Placebo effect (Δ control – placebo: green) and nocebo effect (Δ nocebo – control: red) based on perceived pain intensity ratings in the test phase. (C) Positive expectations (Δ control – placebo: green) and negative expectations (Δ nocebo – control: red) based on expected pain intensity ratings in the test phase. Expected pain intensity ratings were acquired in the test phase only. Boxplots display the median and interquartile range. Colored circles indicate individual participant means. White circles indicate group means. Gray lines connect each participant’s paired scores within each panel. White circles indicate group means. Asterisks indicate significant differences (\**p* < 0.05, n.s. = not significant). VAS = visual analogue scale.

#### 3.1.2 Placebo and nocebo effects during the test phase

A repeated measures ANOVA for perceived pain intensity ratings during the test phase revealed a significant main effect of *condition* (*F*(2,38) = 30.52, *p* < 0.001), but no main effect of *test phase block* (*F*(2,38) = 0.78, *p* = 0.457) and no *test phase block* × *condition* interaction (*F*(4,76) = 0.21, *p* = 0.867). Post hoc *t*-tests showed a significant placebo effect indicated by lower perceived pain intensity ratings in the placebo compared to the control condition (mean difference = -11.05; *t*(19) = -5.90, *p* < 0.001, 95% CI [-14.97, -7.13]) and a significant nocebo effect indicated by higher perceived pain intensity ratings in the nocebo compared to the control condition (mean difference = 5.88; *t*(19) = 3.69, *p* = 0.002, 95% CI [2.54, 9.21]. Within each test phase block, placebo and nocebo effects were also significant (*p*s ≤ 0.049; **Figure 2A; Supplementary Table 3)**. While conditioned at a similar strength, the placebo effect was stronger than the nocebo effect across the test phase (*t*(19) = 2.72, *p* = 0.014, 95% CI [1.19, 9.16]; **Figure 3B**).

#### 3.1.3 Positive and negative expectations during the test phase

A repeated measures ANOVA for expected pain intensity ratings during the test phase revealed a significant main effect of *condition* (*F*(2,38) = 85.08, *p* < 0.001) and a significant *test phase block* × *condition* interaction (*F*(4,76) = 7.05, *p* < 0.001), but no main effect of *test phase block* (*F*(2,38) = 1.11, *p* = 0.333). Post hoc *t*-tests showed significant positive expectations indicated by lower expected pain intensity ratings in the placebo compared to the control condition (mean difference = -18.55; *t*(19) = -8.35, *p* < 0.001, 95% CI [-23.20, -13.90]) and significant negative expectations indicated by higher expected pain intensity ratings in the nocebo compared to the control condition (mean difference = 15.19; *t*(19) = 8.49, *p* < 0.001, 95% CI [11.44, 18.94]). Within each test phase block, positive and negative expectations were also significant (*p*s < 0.001; **Figure 2A**; **Supplementary Table 4**). Across the test phase, the magnitudes of the positive and negative expectations did not differ (*t*(19) = 1.62, *p* = 0.121, 95% CI [-0.97, 7.68]; **Figure 3C**). With regard to the significant *test phase block* x *condition* interaction, expected pain intensity ratings decreased significantly from Test 1 to Test 2 in the nocebo condition (mean difference = 8.46, *t*(19) = 3.78, *p* = 0.011, 95% CI [3.77, 13.14]), whereas no other within-condition block-to-block changes were significant (*p*s ≥ 0.491; **Supplementary Table 5**).

### 3.2 EEG results

#### 3.2.1 Differences in EEG activity between conditions

A cluster-corrected dependent samples *t*-test revealed a significant positive cluster characterized by increased theta, alpha and beta power in the placebo compared to the control condition during pain delivery (*p* = 0.006, 4.03-19.04 Hz, 0.10 to 0.95 sec relative to heat stimulus onset; **Figure 4A**). While most electrodes were involved in the cluster, the effect was most pronounced at central and parietal regions (**Figure 4B**).

**Figure 4.**
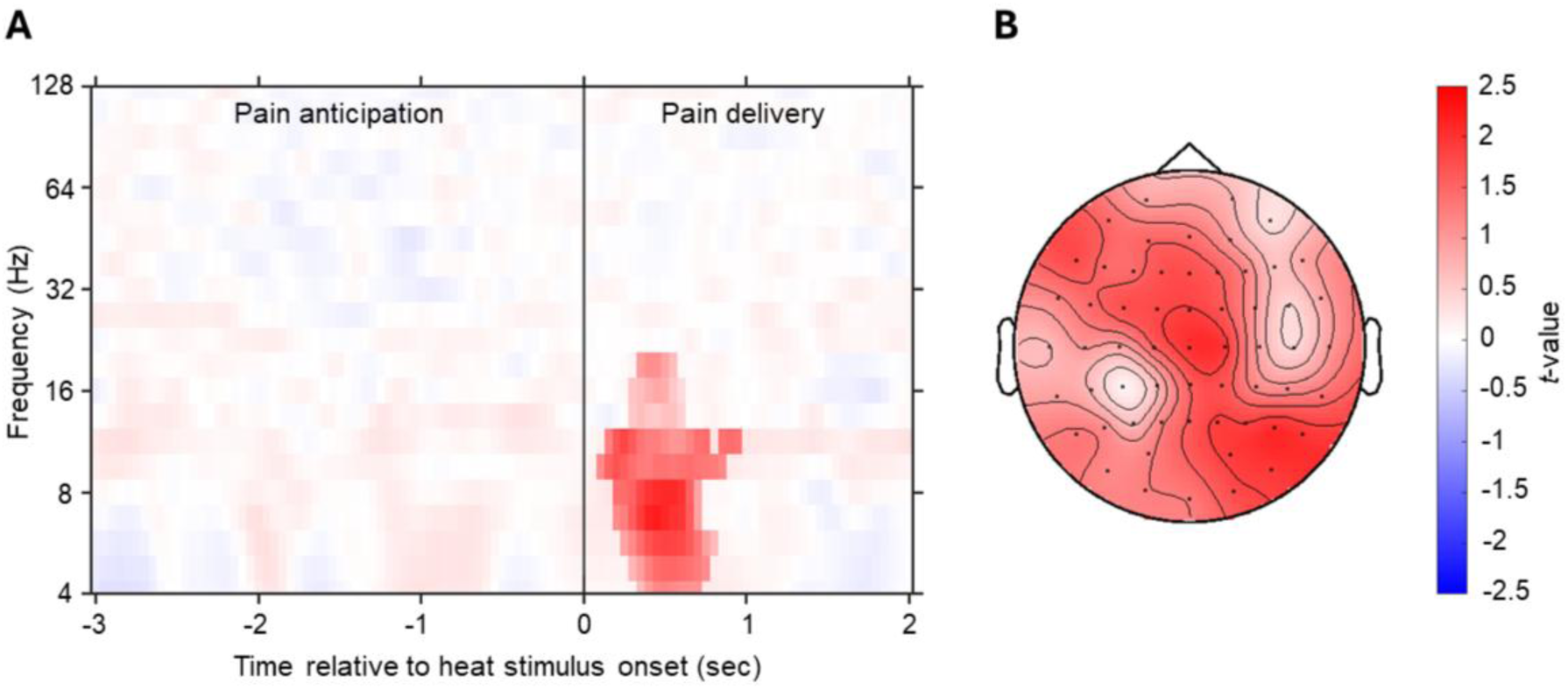
EEG activity in the placebo compared to the control condition. A) Time-frequency plot during pain anticipation (-3 to 0 sec relative to heat stimulus onset) and pain delivery (0 to 2 sec relative to heat stimulus onset) including the significant cluster. *T*-values were averaged across electrodes. Non-significant time-frequency points are masked. B) Topography of the cluster. *T*-values were averaged per electrode across all time-frequency points within the cluster.

A cluster-corrected dependent samples *t*-test revealed a significant positive cluster characterized by increased theta, alpha and beta power in the nocebo compared to the control condition at the beginning of the pain anticipation period (*p* = 0.050, 4.03-22.58 Hz, -2.35 to -1.70 sec relative to heat stimulus onset; **Figure 5A**). While most electrodes were involved in the cluster, the effect was most dominant at centroparietal and temporal regions (**Figure 5B**). A second positive cluster was significant in the nocebo compared to the control condition, characterized by increased theta, alpha and beta power directly before heat stimulus onset and during pain delivery (*p* = 0.006, 4.03-15.99 Hz, -0.75 to 0.85 sec relative to heat stimulus onset; **Figure 5A**). While most electrodes were involved in the cluster, the effect was most pronounced at frontocentral and parieto-occipital regions (**Figure 5B**).

**Figure 5.**
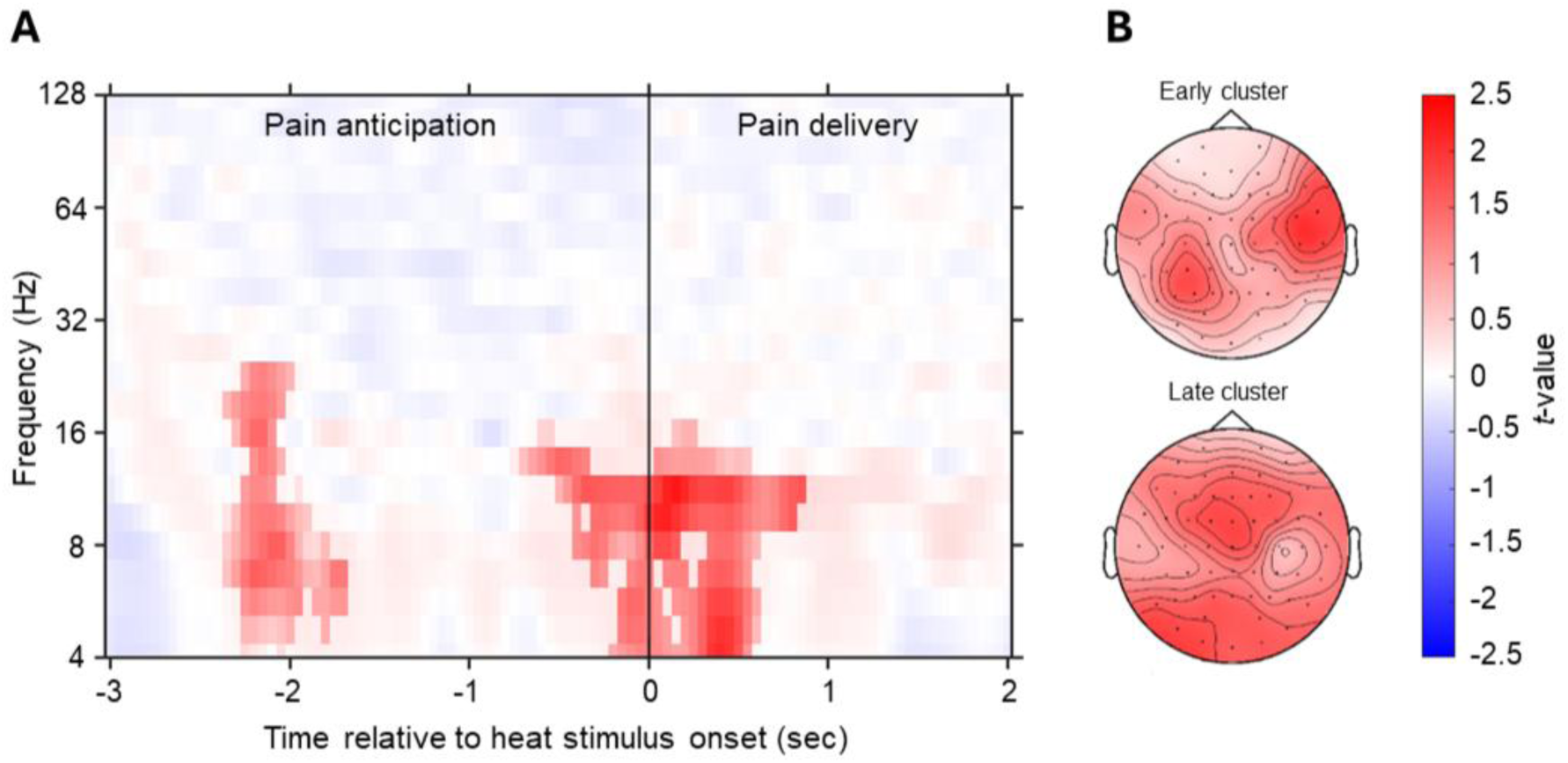
EEG activity in the nocebo compared to the control condition. A) Time-frequency plot during pain anticipation (-3 to 0 sec relative to heat stimulus onset) and pain delivery (0 to 2 sec relative to heat stimulus onset) including the significant clusters. *T*-values were averaged across electrodes. Non-significant time-frequency points are masked. B) Topography of the early cluster (i.e., at the beginning of the pain anticipation period) and late cluster (i.e., directly before heat stimulus onset and during pain delivery). *T*-values were averaged per electrode across all time-frequency points within the cluster.

No significant clusters were found in the cluster-corrected dependent samples *t*-test that compared the nocebo with the placebo condition (*p* ≥ 0.588).

A cluster-corrected dependent samples *t*-test revealed a significant positive cluster in the shared expectation (placebo and nocebo) compared to the control condition (*p* = 0.006, 4.03-19.04 Hz, -0.75 to 1 sec relative to heat stimulus onset; **Figure 6A**). Theta, alpha and beta power was increased in both the placebo and nocebo compared to the control condition directly before heat stimulus onset and during pain delivery. While most electrodes were involved in the cluster, the effect was most pronounced at frontocentral and parieto-occipital regions (**Figure 6B**).

**Figure 6.**
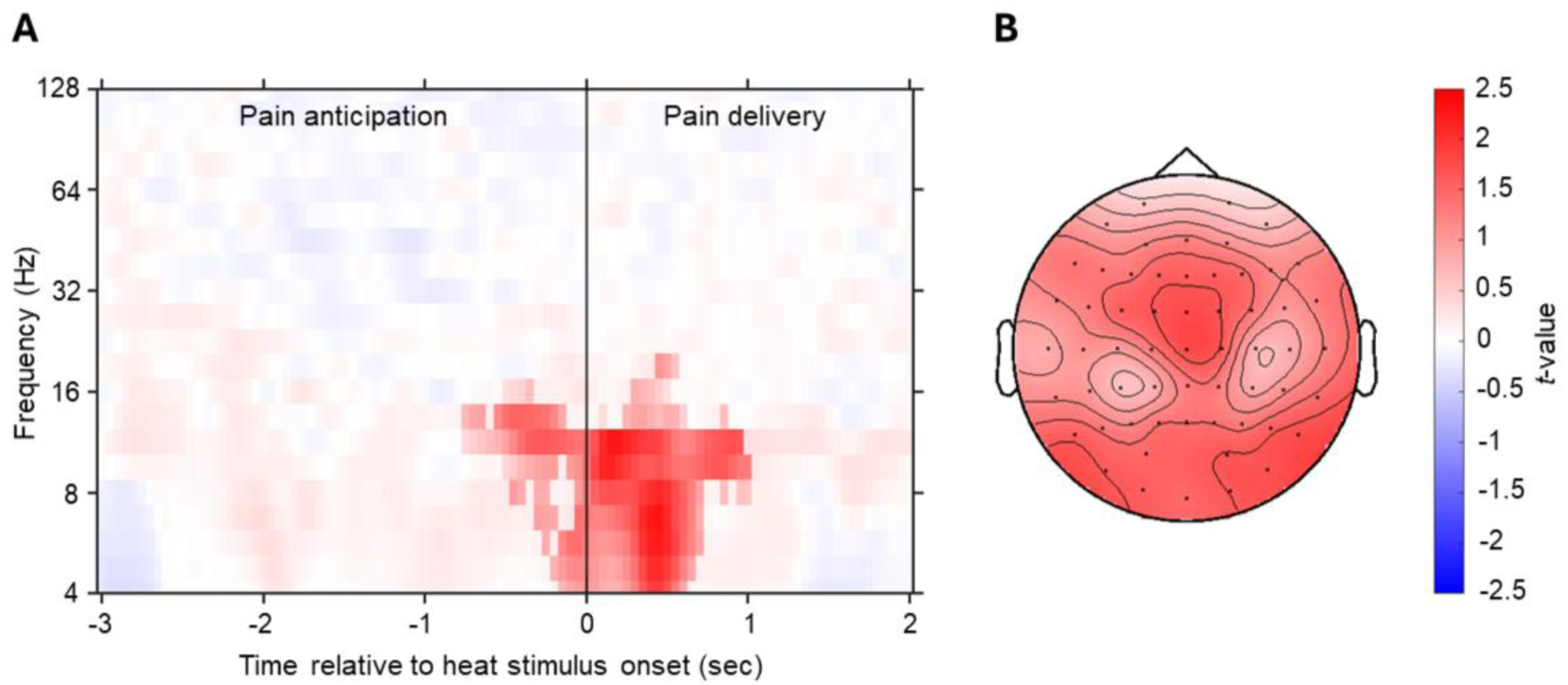
EEG activity in the shared expectation (placebo and nocebo) compared to the control condition. A) Time-frequency plot during pain anticipation (-3 to 0 sec relative to heat stimulus onset) and pain delivery (0 to 2 sec relative to heat stimulus onset) including the significant cluster. *T*-values were averaged across electrodes. Non-significant time-frequency points are masked. B) Topography of the cluster. *T*-values were averaged per electrode across all time-frequency points within the cluster.

##### 3.2.1.1 Correlations of the clusters with psychological questionnaires

To gain additional insights in the processes related to the EEG clusters, an exploratory post hoc analysis was performed to examine Spearman correlations between power in the clusters from the dependent samples *t*-tests and psychological questionnaires that assessed individual traits and states (see Supplementary Materials for analysis details). Power in the placebo compared to the control condition cluster during pain delivery was not significantly associated with any of the questionnaires (*p* ≥ 0.123; **Supplementary Table 6**). More power in the nocebo compared to the control condition cluster at the beginning of the pain anticipation period was associated with a lower pain catastrophizing (PCS) total score (*rho* = -0.498, *p* = 0.025). No other associations were significant for this cluster (*p* ≥ 0.084; **Supplementary Table 7**). Power in the nocebo compared to the control condition cluster directly before heat stimulus onset and during pain delivery was significantly positively associated with FPQ-III fear of medical pain (*rho* = 0.480, *p* = 0.032) and fear of minor pain (*rho* = 0.464, *p* = 0.039), BFI-10 neuroticism (*rho* = 0.536, *p* = 0.015), STADI trait depression (*rho* = 0.470, *p* = 0.037) and the STADI state anxiety and depression total score before the test phase (*rho* = 0.538, *p* = 0.014), which was driven by state depression (*rho* = 0.500, *p* = 0.025). No other associations were significant for this cluster (*p* ≥ 0.064; **Supplementary Table 8**). More power in the shared expectation (placebo and nocebo) compared to the control condition cluster directly before heat stimulus onset and during pain delivery was associated with a higher BFI-10 neuroticism score (*rho* = 0.569, *p* = 0.009). No other associations were significant for this cluster (*p* ≥ 0.099; **Supplementary Table 9**).

#### 3.2.2 EEG activity as a function of perceived pain intensity differences

The linear regression revealed no significant clusters for the association between the behavioral placebo effect (control – placebo perceived pain intensity rating difference) and EEG activity in the placebo compared to the control condition (*p* ≥ 0.304). No significant clusters were found for the association between the behavioral nocebo effect (nocebo – control perceived pain intensity rating difference) and EEG activity in the nocebo compared to the control condition either (*p* ≥ 0.097). With regard to the association between the nocebo - placebo perceived pain intensity rating difference and EEG activity in the nocebo compared to the placebo condition, the linear regression revealed one significant positive cluster (*p* = 0.004, 26.86-128.05 Hz, - 1.05 to 2 sec relative to heat stimulus onset; **Figure 7A**). Participants who reported more pain in the nocebo than in the placebo condition showed increased gamma power in the nocebo compared to the placebo condition at the end of the pain anticipation period and during pain delivery. While most electrodes were involved in the cluster, the effect was most pronounced at centroparietal regions (**Figure 7B**).

**Figure 7.**
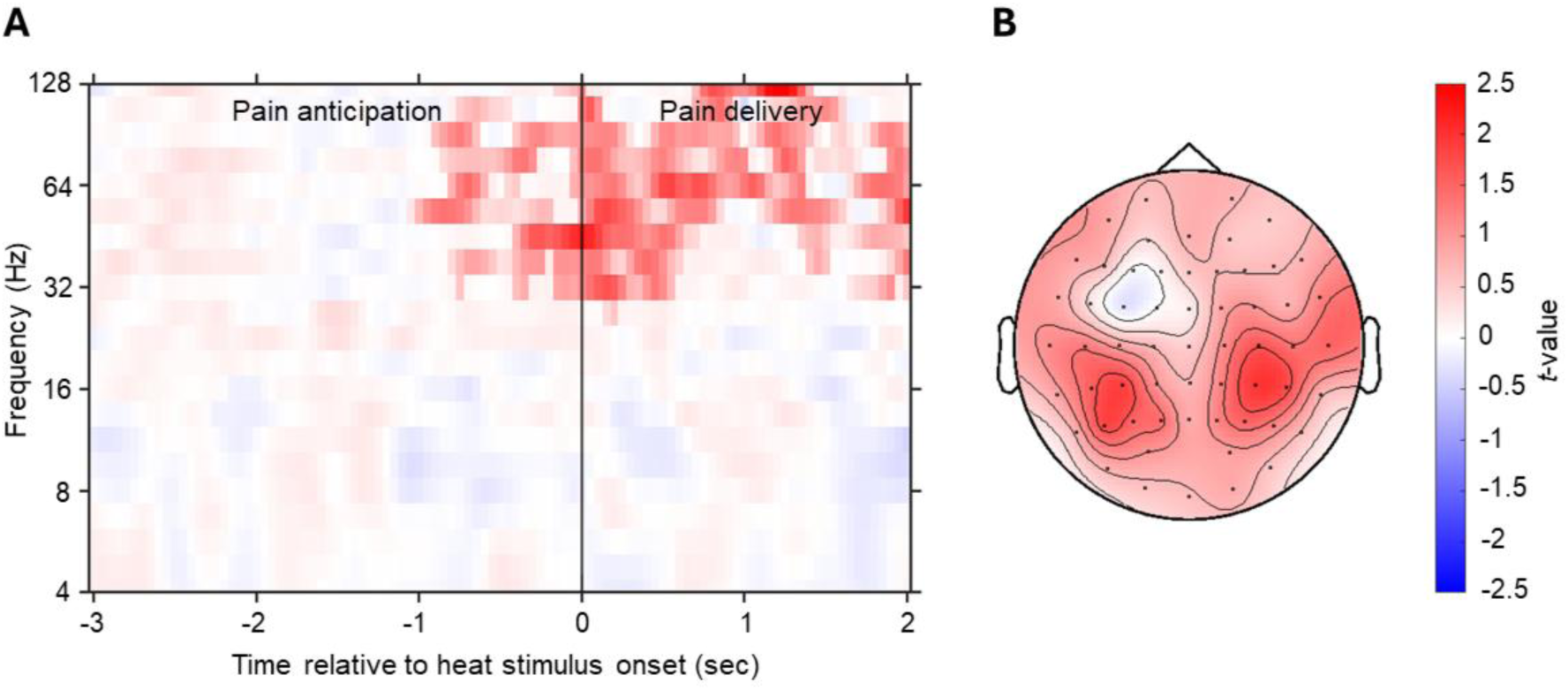
Positive association between the nocebo - placebo perceived pain intensity rating difference and EEG activity in the nocebo compared to the placebo condition. A) Time-frequency plot during pain anticipation (-3 to 0 sec relative to heat stimulus onset) and pain delivery (0 to 2 sec relative to heat stimulus onset) including the significant cluster. *T*-values were averaged across electrodes. Non-significant time-frequency points are masked. B) Topography of the cluster. *T*-values were averaged per electrode across all time-frequency points within the cluster.

## 4. Discussion

The primary aim of this feasibility study was to assess adaptations made to an established placebo-nocebo conditioning pain paradigm (German Clinical Trial Register, 2023; Kunkel et al., 2025) in the context of EEG. Results showed placebo and nocebo effects, as in the other studies that used previous versions of the paradigm (German Clinical Trial Register, 2023; Kunkel et al., 2025). The EEG-adapted version of the paradigm, with a substantially shorter duration of the heat stimulus (i.e., from 20 to 2 sec) and more trials in the test phase, was thus effective. Moreover, based on the trial-by-trial expected pain intensity ratings that were implemented in the paradigm in this study, positive and negative expectations were demonstrated. That is, a lower pain intensity was expected in the placebo compared to the control condition, and a higher pain intensity was expected in the nocebo compared to the control condition. Placebo and nocebo effects remained stable across the test phase, as did positive expectations, although negative expectations decreased from the first to the second test phase block. Notably, this decrease in negative expectations was not accompanied by a reduction of the nocebo effect across test phase blocks. This study found a stronger placebo than nocebo effect, whereas a study that used a previous version of the paradigm reported a stronger nocebo than placebo effect (Kunkel et al., 2025). Importantly, pain relief and pain worsening were conditioned at a similar strength. Arguably, our small sample size as well as differences in context (e.g., EEG) may account for the opposite finding.

The secondary aim of the study was to assess EEG correlates of placebo hypoalgesia and nocebo hyperalgesia during the anticipation and delivery of experimental heat pain in healthy adult volunteers. Partly in line with our hypothesis, theta, alpha and beta power were increased in the placebo compared to the control condition during pain delivery, and in the nocebo compared to the control condition during pain anticipation and delivery. The increase in theta, alpha and beta power directly before heat stimulus onset and during pain delivery was shared by the placebo and the nocebo conditions. Furthermore, in line with our hypothesis, power in the gamma-band was increased in the nocebo compared to the placebo condition as a function of a higher perceived pain intensity. However, no associations were found between EEG activity and the behavioral placebo and nocebo effect.

The increases in alpha and beta power in the placebo and nocebo conditions during pain anticipation and delivery support the idea that alpha- and beta-band activity are related to top-down processing in the context of pain (Ploner et al., 2017). Interestingly, and in contrast to our hypothesis, theta power was also increased in these conditions, which points to a role for theta-band activity in top-down pain processing as well (Tiemann et al., 2015). During pain anticipation in particular, an increase in alpha and beta power have been related to the processing of pain stimulus intensity expectations (Strube et al., 2021). Furthermore, theta- and alpha-band activity have, respectively, been linked to emotional appraisal (Knyazev, 2007) and self-referential thought (Knyazev, 2013; Knyazev et al., 2011), and these processes underlie placebo and nocebo effects (Ashar et al., 2017; Colloca, 2024). In the context of placebo hypoalgesia and nocebo hyperalgesia, the increase in theta and alpha power may therefore reflect such cognitive-affective interpretation processes, probably particularly during pain delivery. The increase in theta, alpha and beta power directly before heat stimulus onset and during pain delivery was shared by the placebo and nocebo conditions, indicating a valence-independent process. Arguably, this cluster may reflect expectation processing before heat stimulus onset, which transitions into evaluation of the heat pain stimulus in the light of those expectations during pain delivery.

In contrast, the increase in theta, alpha and beta power at the beginning of the pain anticipation period was present only in the nocebo (compared to the control) condition, suggesting enhanced anticipatory cognitive-affective processing of negative pain-related expectations. Support for this idea comes from a study that reported a more negative stimulus-preceding negativity (a measure of top-down processing) during pain anticipation in the nocebo compared to the control condition (Blythe et al., 2023). Moreover, increased theta, alpha and beta power was found during the anticipation of higher expected pain intensities (Strube et al., 2021), as well as in a nocebo compared to placebo condition (Wolf et al., 2024). Additionally, changes in theta power were reported during the anticipation of a painful compared to non-painful stimulus, which originated from the insula (Taesler & Rose, 2016). Another study found that activity in the anterior insula during pain anticipation for both positive (placebo) and negative (nocebo) expectations (i.e., valence-independent) was related to lower activity in the theta to lower gamma frequency band (Wittkamp et al., 2024). During pain anticipation, the anterior insula is involved in expectation generation and signaling salience of the expected pain level, for more painful (Taesler & Rose, 2016; Wiech et al., 2010) and valence-independent (Wittkamp et al., 2024) expectations. It should be noted that salience processing goes beyond bottom-up processing of external stimulus properties, and also involves assessment of homeostatic relevance in the light of cognitive, emotional and motivational states, for which particularly the anterior insula is known (Borsook et al., 2013; Craig, 2003, 2009). This role of the anterior insula and the association with anticipatory theta-, alpha- and beta-band activity may suggest that the cluster in the nocebo condition at the beginning of the pain anticipation period reflects increased salience processing for negative pain expectations. However, in that case, a difference in EEG activity may also have been expected during pain anticipation in the nocebo compared to the placebo condition, which was not observed. A possibility for this absent finding is that pain anticipation in the placebo condition also elicits some form of salience.

The negative association between power in the cluster at the beginning of the pain anticipation period in the nocebo versus control condition and the pain catastrophizing total score may seem counterintuitive. This finding may be interpreted as a difficulty to dissociate the potential threat between the two conditions. It should be noted, however, that the differences in perceived and expected pain intensity ratings between the two conditions are not in line with this interpretation. The cluster directly before heat stimulus onset and during pain delivery in the nocebo versus control condition was positively associated with various negative traits and states. The positive associations with fear of medical and minor pain, neuroticism, trait depression, and state anxiety and depression concur with positive associations reported in the literature between nocebo hyperalgesia and these negative traits and states (Aslaksen & Lyby, 2015; Corsi & Colloca, 2017; Karacaoglu et al., 2023; Kern et al., 2020; Thomaidou et al., 2021b). Notably, neuroticism was positively related to the shared expectation (i.e., placebo and nocebo) versus control condition cluster, in line with positive associations reported between neuroticism and placebo as well as nocebo effects (Fischer et al., 2026; Rosén et al., 2017). Taken together, these results suggest that, in the nocebo condition, particularly EEG activity directly before heat stimulus onset and during pain delivery is related to negative emotional traits and states, in contrast to early pain anticipation. This idea is in line with evidence showing that fear of high pain amplified nocebo hyperalgesia when high pain was delivered but not when it was anticipated (Thomaidou et al., 2021b). Arguably, EEG activity during pain delivery in the nocebo condition may be more related to affective processes, which may be the evaluation of pain in the light of negative emotions. Speculatively, EEG activity at the beginning of the pain anticipation period in the nocebo condition, on the other hand, may be more related to (more cognitive) saliency processes, as also suggested in the previous paragraph. It should be noted that the correlations between power in EEG clusters and psychological constructs were exploratory and therefore not corrected for multiple testing. Future research is needed to further test the hypothesis of a potential dissociation between more cognitive versus affective processes in the context of negative pain-related expectations during pain anticipation and delivery, respectively.

Based on the association with perceived pain intensity, our results suggest that gamma-band activity, in addition to stimulus intensity-driven bottom-up processing (Ploner et al., 2017; Tiemann et al., 2015; Strube et al., 2021), also reflects sensory processing modulated by expectations when actual stimulus intensities are the same. In line with this idea, the increase in gamma power was dominant at centroparietal regions, in keeping with previous studies that demonstrated an effect of perceived pain intensity at the primary somatosensory cortex (Gross et al., 2007; Zhang et al., 2012). However, no differences were found in both the placebo and the nocebo condition compared to the control condition as a function of differences in perceived pain intensity. Differences in perceived pain intensity between those conditions may not have been strong enough to affect EEG activity in our relatively small sample.

The small sample size is a limitation of the study. A power analysis was not performed a priori to determine the sample size, given that the study was primarily designed as a feasibility study. A small sample size increases the possibility of type I and type II errors (Button et al., 2013; Christley, 2010), and therefore, our results should be interpreted with caution.

In conclusion, feasibility of the EEG-compatible placebo-nocebo conditioning pain paradigm was demonstrated. EEG results are in line with previous findings that, in the context of placebo hypoalgesia and nocebo hyperalgesia, link theta-, alpha- and beta-band activity to top-down processing, and gamma-band activity to expectation-modulated sensory processing.

## Declaration of interest

The authors have no conflicts of interest to declare.

## Funding

The work is funded by the Deutsche Forschungsgemeinschaft (DFG, German Research Foundation) - Project-ID 422744262 - TRR 289.

## CrediT authorship contribution statement

**Antonia Baumgarten**: Conceptualization, Investigation, Methodology, Project administration, Software, Visualization, Writing – original draft. **Frederik Schlitt-Nguyen**: Conceptualization, Methodology, Supervision, Writing – review & editing. **Helena Hartmann**: Methodology, Writing – review & editing. **Katharina Schmidt**: Writing – review & editing. **Michael Rose**: Conceptualization, Methodology, Software, Writing – review & editing. **Dagmar Timmann**: Conceptualization, Funding acquisition, Methodology, Project administration, Supervision, Writing – review & editing. **Enzo Nio**: Conceptualization, Data curation, Formal analysis, Methodology, Software, Supervision, Visualization, Writing – original draft. **Eline S. Kruithof**: Conceptualization, Formal analysis, Methodology, Software, Supervision, Visualization, Writing – original draft.

## Acknowledgements

The authors thank Thomas Ernst for the technical support and Greta Wippich for the illustration of the experimental setup and electrode and thermode position.

## Data statement

All data are available upon reasonable request to the corresponding author.

## Notes

### Competing Interest Statement

The authors have declared no competing interest.

