## Supplementary materials for "Electrophysiological correlates of placebo hypoalgesia and nocebo hyperalgesia during the anticipation and delivery of pain"

Questionnaire information and descriptive statistics

**Supplementary Table 1**. Psychological questionnaires. For all questionnaires, the German version was used. A higher score indicated a higher level of the respective measure. Time point (T) indicates when the questionnaire was administered: T1 = baseline, T2 = before the practice trials, T3 = before the test phase, T4 =after the test phase. BFI-10 = Big Five Inventory-10, FPQ-III = Fear of Pain Questionnaire-III, ISAQ = Interoceptive Awareness and Sensitivity Questionnaire, NA = not applicable, PCS = Pain Catastrophizing Scale, PSS-10 = Perceived Stress Scale-10, SSAS = Somatosensory Amplification Scale, STADI = State-Trait-Anxiety-Depression Inventory.

| **Questionnaire** | **Subscales and/or total score (number of items)** | **Response scale** | **Time point** | **Citation German version** | **Citation English version (if applicable)** |
| --- | --- | --- | --- | --- | --- |
| Big Five Inventory-10 (BFI-10) | Extraversion (2), agreeableness (2), conscientiousness (2), neuroticism (2), openness (2) | 1 (‘disagree strongly’) to 5 (‘agree strongly’) | T1 | Rammstedt & John (2007) | Rammstedt & John (2007) |
| Fear of Pain Questionnaire III (FPQ-III) | Fear of severe pain (10), fear of minor pain (10), fear of medical pain (10), total score (30) | 1 (‘not at all’) to 5 (‘extreme’) | T1 | Baum et al. (2013) | McNeil & Rainwater (1998) |
| Generic rating scale for previous treatment experiences, treatment expectations, and treatment effects (GEEE) | Previous treatment experiences with the pain-relieving electrical stimulation (3) and pain-worsening electrical stimulation (3) | 0 (‘no improvement/worsening/side effects) to 10 (‘largest imaginable improvement/worsening/side effects’) | T1 | Rief et al. (2021) | NA |
|  | Treatment expectations for the pain-relieving electrical stimulation (3) and pain-worsening electrical stimulation (3) | 0 (‘no improvement/worsening/side effects) to 10 (‘largest imaginable improvement/worsening/side effects’) | T2, T3 |  |  |
|  | Experienced treatment effects for the pain-relieving electrical stimulation (3) and pain-worsening electrical stimulation (3) | 0 (‘no improvement/worsening/side effects) to 10 (‘largest imaginable improvement/worsening/side effects’) | T3, T4 |  |  |

Table continued on next page

Supplementary Table 1 (continued)

| **Questionnaire** | **Subscales and/or total score (number of items)** | **Response scale** | **Time point** | **Citation German version** | **Citation English version (if applicable)** |
| --- | --- | --- | --- | --- | --- |
| Interoceptive Sensitivity and Attention Questionnaire (ISAQ) | Attention to unpleasant bodily sensations (4), difficulty disengaging from unpleasant bodily sensations (4), sensitivity to neutral bodily sensations (9) | 1 (‘strongly disagree’) to 5 (‘strongly agree’) | T1 | Pohl et al. (2025) | Bogaerts et al. (2022) |
| Pain Catastrophizing Scale (PCS) | Helplessness (6), magnification (3), rumination (4), total score (13) | 0 (‘not at all’) to 4 (‘all the time’) | T1 | Meyer et al. (2008) | Sullivan et al. (1995) |
| Perceived Stress Scale (PSS-10) | Total score (10) | 0 (‘never’) to 4 (‘very often’) | T1 | Klein et al. (2016) | Cohen et al. (1983) |
| Somatosensory Amplification Scale (SSAS) | Total score (10) | 1 (‘not at all true’) to 5 (‘extremely true’) | T1 | Doering et al. (2015) | Barsky et al. (1988) |
| State-Trait Anxiety-Depression Inventory (STADI) | Trait anxiety (10), trait depression (10), total score (20) | 1 (‘almost never’) to 4 (‘almost always’) | T1 | Laux et al. (2013) | NA |
|  | State anxiety (10), state depression (10), total score (20) | 1 (‘not at all’) to 4 (‘very much’) | T2, T3, T4 |  |  |
| Warmth & competence | Perceived warmth (6) and competence (6) of the main experiment leader | 1 (‘not at all’) to 5 (‘extremely’) | T4 | Seefeld & Rief (2023) | NA |

**Supplementary Table 2**. Descriptive statistics (*M* ± *SD*) of the psychological questionnaires. Time point indicates when the questionnaire was administered: T1 = baseline, T2 = before the practice trials, T3 = before the test phase, T4 =after the test phase. The GEEE at T1 only includes data from one participant, as the other participants had no previous treatment experience. BFI-10 = Big Five Inventory-10, FPQ-III = Fear of Pain Questionnaire-III, GEEE = Generic rating scale for previous treatment experiences, treatment expectations, and treatment effects, ISAQ = Interoceptive Awareness and Sensitivity Questionnaire, NA = not applicable, PCS = Pain Catastrophizing Scale, PSS-10 = Perceived Stress Scale-10, SSAS = Somatosensory Amplification Scale, STADI = State-Trait-Anxiety-Depression Inventory.

| **Questionnaire** | **Time point** | **Subscales and/or total score** | **Subquestions (GEEE only)** | ***M* ± *SD*** |
| --- | --- | --- | --- | --- |
| BFI-10 |  |  |  |  |
|  | T1 |  |  |  |
|  |  | Agreeableness | NA | 3.3 ± 1.03 |
|  |  | Conscientiousness | NA | 3.38 ± 0.84 |
|  |  | Extraversion | NA | 3.1 ± 1.07 |
|  |  | Neuroticism | NA | 2.88 ± 1.07 |
|  |  | Openness | NA | 3.55 ± 1.06 |
| FPQ-III |  |  |  |  |
|  | T1 |  |  |  |
|  |  | Medical pain | NA | 22.05 ± 7.83 |
|  |  | Minor pain | NA | 16.55 ± 5.44 |
|  |  | Severe pain | NA | 32.5 ± 6.13 |
|  |  | Total score | NA | 71.1 ± 16.29 |
| GEEE |  |  |  |  |
|  | T1 (*n* = 1) |  |  |  |
|  |  | Pain-relieving electrical stimulation |  |  |
|  |  |  | Previously experienced improvement | 7 ± 0 |
|  |  |  | Previously experienced worsening | 4 ± 0 |
|  |  |  | Previously experienced side effects | 0 ± 0 |
|  |  | Pain-worsening electrical stimulation | |  |
|  |  |  | Previously experienced improvement | 5 ± 0 |
|  |  |  | Previously experienced worsening | 5 ± 0 |
|  |  |  | Previously experienced side effects | 2 ± 0 |
|  | T2 | | |  |
|  |  | Pain-relieving electrical stimulation | |  |
|  |  |  | Expected improvement | 5.45 ± 2.06 |
|  |  |  | Expected worsening | 1.3 ± 2.43 |
|  |  |  | Expected side effects | 0.9 ± 1.07 |
|  |  | Pain-worsening electrical stimulation | |  |
|  |  |  | Expected improvement | 1.15 ± 1.87 |
|  |  |  | Expected worsening | 5.45 ± 2.52 |
|  |  |  | Expected side effects | 3.5 ± 2.86 |

Table continued on next page

Supplementary Table 2 (continued)

| **Questionnaire** | **Time point** | **Subscales and/or total score** | **Subquestions (GEEE only)** | ***M* ± *SD*** |
| --- | --- | --- | --- | --- |
| GEEE |  |  |  |  |
|  | T3 | | |  |
|  |  | Pain-relieving electrical stimulation | |  |
|  |  |  | Experienced improvement | 6.15 ± 2.11 |
|  |  |  | Experienced worsening | 1.55 ± 2.78 |
|  |  |  | Experienced side effects | 1.15 ± 2.25 |
|  |  |  | Expected improvement | 6.55 ± 2.16 |
|  |  |  | Expected worsening | 0.45 ± 1 |
|  |  |  | Expected side effects | 0.55 ± 1.05 |
|  |  | Pain-worsening electrical stimulation | |  |
|  |  |  | Experienced improvement | 0.95 ± 1.82 |
|  |  |  | Experienced worsening | 6.55 ± 2.42 |
|  |  |  | Experienced side effects | 3.2 ± 3.09 |
|  |  |  | Expected improvement | 0.7 ± 1.69 |
|  |  |  | Expected worsening | 6.05 ± 2.37 |
|  |  |  | Expected side effects | 3.4 ± 3.1 |
|  | T4 | | |  |
|  |  | Pain-relieving electrical stimulation | |  |
|  |  |  | Experienced improvement | 2.7 ± 1.95 |
|  |  |  | Experienced worsening | 1.1 ± 1.52 |
|  |  |  | Experienced side effects | 1.4 ± 2.06 |
|  |  | Pain-worsening electrical stimulation | |  |
|  |  |  | Experienced improvement | 1 ± 1.81 |
|  |  |  | Experienced worsening | 3.3 ± 2.87 |
|  |  |  | Experienced side effects | 2.35 ± 2.7 |
| ISAQ |  |  |  |  |
|  | T1 |  |  |  |
|  |  | Attention | NA | 12.75 ± 3.18 |
|  |  | Disengagement | NA | 8.75 ± 3.23 |
|  |  | Sensitivity | NA | 29.25 ± 5.68 |
| PCS |  |  |  |  |
|  | T1 |  |  |  |
|  |  | Helplessness | NA | 7.4 ± 5.65 |
|  |  | Magnification | NA | 4.6 ± 3.03 |
|  |  | Rumination | NA | 8.1 ± 4.52 |
|  |  | Total score | NA | 20.1 ± 11.77 |
| PSS-10 |  |  |  |  |
|  | T1 |  |  |  |
|  |  | Total score | NA | 18.3 ± 6.67 |
| SSAS |  |  |  |  |
|  | T1 |  |  |  |
|  |  | Total score | NA | 27.9 ± 4.99 |
| STADI |  |  |  |  |
|  | T1 |  |  |  |
|  |  | Trait anxiety | NA | 20.8 ± 6.66 |
|  |  | Trait depression | NA | 18.1 ± 5.07 |
|  |  | Total score | NA | 38.9 ± 10.92 |
|  | T2 |  |  |  |
|  |  | State anxiety | NA | 14.95 ± 4.12 |
|  |  | State depression | NA | 18.2 ± 2.86 |
|  |  | Total score | NA | 33.15 ± 5.7 |
|  | T3 |  |  |  |
|  |  | State anxiety | NA | 15.75 ± 6.72 |
|  |  | State depression | NA | 19.05 ± 4.44 |
|  |  | Total score | NA | 34.8 ± 9.77 |
|  | T4 |  |  |  |
|  |  | State anxiety | NA | 14.95 ± 6.52 |
|  |  | State depression | NA | 18.75 ± 4.41 |
|  |  | Total score | NA | 33.7 ± 9.63 |
| Warmth & competence |  |  |  |  |
|  | T4 |  |  |  |
|  |  | Warmth of the main experiment leader | NA | 3.66 ± 0.45 |
|  |  | Competence of the main experiment leader | NA | 3.82 ± 0.28 |

Placebo and nocebo effects and positive and negative expectations

**Supplementary Table 3.** Placebo (placebo versus control) and nocebo (nocebo versus control) effects per test phase block (Test 1, Test 2, Test 3). *df* = degrees of freedom, CI = confidence interval, MD = mean difference, *p*(Holm) = Holm-corrected *p*-value.

| **Test phase block** | **Contrast** | **MD** | ***t*** | ***df*** | ***p*(Holm)** | **95% CI** |
| --- | --- | --- | --- | --- | --- | --- |
| Test 1 | Placebo effect | -11.27 | -4.97 | 19 | < 0.001 | [-16.00, -6.53] |
| Test 1 | Nocebo effect | 5.51 | 2.31 | 19 | 0.049 | [0.52, 10.50] |
| Test 2 | Placebo effect | -9.84 | -3.96 | 19 | 0.003 | [-15.03, -4.64] |
| Test 2 | Nocebo effect | 6.70 | 4.59 | 19 | < 0.001 | [3.65, 9.75] |
| Test 3 | Placebo effect | -12.04 | -4.71 | 19 | < 0.001 | [-17.40, -6.69] |
| Test 3 | Nocebo effect | 5.42 | 2.44 | 19 | 0.049 | [0.78, 10.06] |

**Supplementary Table 4.** Positive (placebo versus control) and negative (nocebo versus control) expectations per test phase block (Test 1, Test 2, Test 3). *df* = degrees of freedom, CI = confidence interval, MD = mean difference, *p*(Holm) = Holm-corrected *p*-value.

| **Test phase block** | **Contrast** | **MD** | ***t*** | ***df*** | ***p*(Holm)** | **95% CI** |
| --- | --- | --- | --- | --- | --- | --- |
| Test 1 | Positive expectations | 20.21 | 7.62 | 19 | < 0.001 | [-25.76, -14.66] |
| Test 1 | Negative expectations | 19.53 | 8.99 | 19 | < 0.001 | [14.99, 24.07] |
| Test 2 | Positive expectations | 17.59 | 8.47 | 19 | < 0.001 | [-21.94, -13.24] |
| Test 2 | Negative expectations | 11.63 | 5.02 | 19 | < 0.001 | [6.78, 16.47] |
| Test 3 | Positive expectations | 17.84 | 6.77 | 19 | < 0.001 | [-23.35, -12.33] |
| Test 3 | Negative expectations | 14.42 | 6.83 | 19 | < 0.001 | [10.00, 18.83] |

**Supplementary Table 5.** Differences in expected pain intensity ratings between test phase blocks (Test 1, Test 2, Test 3) within each condition, following up the significant *test phase block* × *condition* interaction. *df* = degrees of freedom, CI = 95% confidence interval, MD = mean difference, *p*(Holm) = Holm-corrected *p*-value. Significant comparisons are indicated in bold.

| **Condition** | **Contrast** | **MD** | ***t*** | ***df*** | ***p*(Holm)** | **95% CI** |
| --- | --- | --- | --- | --- | --- | --- |
| Placebo | Test 1 vs Test 2 | -2.07 | -1.17 | 19 | 1.000 | [-5.78, 1.64] |
| Placebo | Test 1 vs Test 3 | -2.47 | -1.45 | 19 | 1.000 | [-6.05, 1.11] |
| Placebo | Test 2 vs Test 3 | -0.4 | -0.37 | 19 | 1.000 | [-2.67, 1.87] |
| Control | Test 1 vs Test 2 | 0.55 | 0.22 | 19 | 1.000 | [-4.57, 5.67] |
| Control | Test 1 vs Test 3 | -0.1 | -0.06 | 19 | 1.000 | [-3.88, 3.68] |
| Control | Test 2 vs Test 3 | -0.65 | -0.41 | 19 | 1.000 | [-3.93, 2.63] |
| **Nocebo** | **Test 1 vs Test 2** | **8.46** | **3.78** | **19** | **0.011** | **[3.77, 13.14]** |
| Nocebo | Test 1 vs Test 3 | 5.01 | 1.99 | 19 | 0.491 | [-0.26, 10.29] |
| Nocebo | Test 2 vs Test 3 | -3.44 | -1.48 | 19 | 1.000 | [-8.29, 1.41] |

EEG

Post hoc exploratory analysis association EEG clusters with psychological questionnaires

To gain additional insights in the processes related to the EEG clusters, we computed Spearman correlations between power in the clusters from the dependent samples *t*-tests and psychological questionnaires that assessed individual traits and states. Average power in each cluster was calculated for each participant by computing the difference in power between the respective conditions. Correlations were analyzed with the BFI-10 agreeableness subscale, BFI-10 conscientiousness subscale, BFI-10 extraversion subscale, BFI-10 neuroticism subscale, BFI-10 openness subscale, FPQ-III medical pain subscale, FPQ-III minor pain subscale, FPQ-III severe pain subscale, FPQ-III total score, ISAQ attention subscale, ISAQ disengagement subscale, ISAQ sensitivity subscale, PCS helplessness subscale, PCS magnification subscale, PCS rumination subscale, PCS total score, PSS-10, SSAS, STADI trait anxiety subscale, STADI trait depression subscale, STADI trait total score, STADI state anxiety subscale at T3 (before the test phase), STADI state depression subscale at T3 (before the test phase) and the STADI state total score at T3 (before the test phase). It should be noted that correlations with the GEEE and Warmth & competence were not analyzed, as these questionnaires do not assess individual traits and states. STADI state total and subscale scores at T3 were used as this was the time point before the test phase.

**Supplementary Table 6.** Spearman correlations between EEG power in the placebo versus control condition cluster and psychological questionnaires that assessed individual traits and states. BFI-10 = Big Five Inventory-10, FPQ-III = Fear of Pain Questionnaire-III, ISAQ = Interoceptive Awareness and Sensitivity Questionnaire, PCS = Pain Catastrophizing Scale, PSS-10 = Perceived Stress Scale-10, SSAS = Somatosensory Amplification Scale, STADI = State-Trait-Anxiety-Depression Inventory. T3 indicates the time point before the test phase.

| **Questionnaire** | ***rho*** | ***p*** |
| --- | --- | --- |
| BFI-10 agreeableness | 0.284 | 0.225 |
| BFI-10 conscientiousness | 0.035 | 0.884 |
| BFI-10 extraversion | 0.049 | 0.838 |
| BFI-10 neuroticism | 0.074 | 0.757 |
| BFI-10 openness | 0.075 | 0.753 |
| FPQ-III medical pain  FPQ-III minor pain  FPQ-III severe pain | -0.103 | 0.667 |
|  | -0.153 | 0.520 |
|  | -0.223 | 0.345 |
| FPQ-III total | -0.171 | 0.471 |
| ISAQ attention | 0.146 | 0.540 |
| ISAQ disengagement | 0.278 | 0.236 |
| ISAQ sensitivity | 0.356 | 0.123 |
| PCS helplessness | -0.153 | 0.520 |
| PCS magnification | -0.316 | 0.175 |
| PCS rumination | -0.098 | 0.683 |
| PCS total | -0.261 | 0.266 |
| PSS-10 | 0.012 | 0.960 |
| SSAS | 0.191 | 0.421 |
| STADI trait anxiety  STADI trait depression | -0.064 | 0.788 |
|  | 0.037 | 0.877 |
| STADI trait total | -0.007 | 0.977 |
| STADI state anxiety at T3  STADI state depression at T3 | 0.167 | 0.481 |
|  | 0.124 | 0.604 |
| STADI state total at T3 | 0.142 | 0.551 |

**Supplementary Table 7.** Spearman correlations of EEG power in the nocebo versus control condition cluster at the beginning of the pain anticipation period with psychological questionnaires that assessed individual traits and states. BFI-10 = Big Five Inventory-10, FPQ-III = Fear of Pain Questionnaire-III, ISAQ = Interoceptive Awareness and Sensitivity Questionnaire, PCS = Pain Catastrophizing Scale, PSS-10 = Perceived Stress Scale-10, SSAS = Somatosensory Amplification Scale, STADI = State-Trait-Anxiety-Depression Inventory. T3 indicates the time point before the test phase. Significant correlations are indicated in bold.

| **Questionnaire** | ***rho*** | ***p*** |
| --- | --- | --- |
| BFI-10 agreeableness | 0.187 | 0.429 |
| BFI-10 conscientiousness | -0.396 | 0.084 |
| BFI-10 extraversion | -0.073 | 0.758 |
| BFI-10 neuroticism | 0.106 | 0.655 |
| BFI-10 openness | 0.083 | 0.728 |
| FPQ-III medical pain  FPQ-III minor pain  FPQ-III severe pain | 0.060 | 0.803 |
|  | 0.067 | 0.778 |
|  | -0.024 | 0.919 |
| FPQ-III total | 0.109 | 0.646 |
| ISAQ attention | 0.095 | 0.690 |
| ISAQ disengagement | -0.069 | 0.774 |
| ISAQ sensitivity | -0.294 | 0.208 |
| PCS helplessness | -0.382 | 0.096 |
| PCS magnification | -0.341 | 0.142 |
| PCS rumination | -0.274 | 0.242 |
| PCS total | **-0.498** | **0.025** |
| PSS-10 | -0.065 | 0.786 |
| SSAS | -0.199 | 0.400 |
| STADI trait anxiety  STADI trait depression | -0.053 | 0.825 |
|  | 0.144 | 0.544 |
| STADI trait total | 0.029 | 0.905 |
| STADI state anxiety at T3  STADI state depression at T3 | -0.095 | 0.692 |
|  | 0.169 | 0.476 |
| STADI state total at T3 | 0.011 | 0.962 |

**Supplementary Table 8.** Spearman correlations of EEG power in the nocebo versus control condition cluster directly before heat stimulus onset and during pain delivery with psychological questionnaires that assessed individual traits and states. BFI-10 = Big Five Inventory-10, FPQ-III = Fear of Pain Questionnaire-III, ISAQ = Interoceptive Awareness and Sensitivity Questionnaire, PCS = Pain Catastrophizing Scale, PSS-10 = Perceived Stress Scale-10, SSAS = Somatosensory Amplification Scale, STADI = State-Trait-Anxiety-Depression Inventory. T3 indicates the time point before the test phase. Significant correlations are indicated in bold.

| **Questionnaire** | ***rho*** | ***p*** |
| --- | --- | --- |
| BFI-10 agreeableness | -0.108 | 0.651 |
| BFI-10 conscientiousness | -0.135 | 0.571 |
| BFI-10 extraversion | 0.017 | 0.943 |
| BFI-10 neuroticism | **0.536** | **0.015** |
| BFI-10 openness | -0.042 | 0.860 |
| FPQ-III medical pain  FPQ-III minor pain  FPQ-III severe pain | **0.480** | **0.032** |
|  | **0.464** | **0.039** |
|  | 0.256 | 0.276 |
| FPQ-III total | 0.416 | 0.068 |
| ISAQ attention | 0.250 | 0.288 |
| ISAQ disengagement | 0.182 | 0.443 |
| ISAQ sensitivity | 0.044 | 0.855 |
| PCS helplessness | 0.135 | 0.569 |
| PCS magnification | 0.071 | 0.768 |
| PCS rumination | -0.043 | 0.857 |
| PCS total | 0.054 | 0.822 |
| PSS-10 | 0.114 | 0.632 |
| SSAS | 0.180 | 0.447 |
| STADI trait anxiety  STADI trait depression | 0.335 | 0.148 |
|  | **0.470** | **0.037** |
| STADI trait total | 0.422 | 0.064 |
| STADI state anxiety at T3  STADI state depression at T3 | 0.397 | 0.083 |
|  | **0.500** | **0.025** |
| STADI state total at T3 | **0.538** | **0.014** |

**Supplementary Table 9.** Spearman correlations between EEG power in the common expectation (placebo and nocebo) versus control condition cluster and psychological questionnaires that assessed individual traits and states. BFI-10 = Big Five Inventory-10, FPQ-III = Fear of Pain Questionnaire-III, ISAQ = Interoceptive Awareness and Sensitivity Questionnaire, PCS = Pain Catastrophizing Scale, PSS-10 = Perceived Stress Scale-10, SSAS = Somatosensory Amplification Scale, STADI = State-Trait-Anxiety-Depression Inventory. T3 indicates the time point before the test phase. Significant correlations are indicated in bold.

| **Questionnaire** | ***rho*** | ***p*** |
| --- | --- | --- |
| BFI-10 agreeableness | 0.130 | 0.585 |
| BFI-10 conscientiousness | -0.052 | 0.828 |
| BFI-10 extraversion | 0.080 | 0.736 |
| BFI-10 neuroticism | **0.569** | **0.009** |
| BFI-10 openness | 0.062 | 0.795 |
| FPQ-III medical pain  FPQ-III minor pain  FPQ-III severe pain | 0.288 | 0.218 |
|  | 0.379 | 0.099 |
|  | 0.198 | 0.403 |
| FPQ-III total | 0.285 | 0.223 |
| ISAQ attention | 0.262 | 0.264 |
| ISAQ disengagement | 0.234 | 0.321 |
| ISAQ sensitivity | 0.207 | 0.382 |
| PCS helplessness | -0.125 | 0.560 |
| PCS magnification | -0.120 | 0.615 |
| PCS rumination | -0.209 | 0.376 |
| PCS total | -0.165 | 0.486 |
| PSS-10 | 0.141 | 0.553 |
| SSAS | 0.244 | 0.299 |
| STADI trait anxiety  STADI trait depression | 0.334 | 0.150 |
|  | 0.329 | 0.157 |
| STADI trait total | 0.374 | 0.104 |
| STADI state anxiety at T3  STADI state depression at T3 | 0.304 | 0.192 |
|  | 0.319 | 0.171 |
| STADI state total at T3 | 0.361 | 0.118 |
